# An Acoustically Controlled Microrobot Modelled on *Spirochete* Bacteria

**DOI:** 10.1101/2023.03.09.531925

**Authors:** Yong Deng, Adrian Paskert, Zhiyuan Zhang, Raphael Wittkowski, Daniel Ahmed

**Affiliations:** Acoustic Robotics Systems Lab (ARSL), Institute of Robotics and Intelligent Systems, ETH Zurich, Rüschlikon, CH-8803, Switzerland; Institut für Theoretische Physik, Center for Soft Nanoscience, Westfälische Wilhelms-Universität Münster, 48149 Münster, Germany

## Abstract

As a next-generation toolkit, microrobots can transform a wide range of fields, including micromanufacturing, electronics, microfluidics, tissue engineering, and medicine. While still in their infancy, acoustically actuated wireless microrobots are becoming increasingly attractive, as acoustic control can generate large propulsive forces, requires relatively simple microrobot design, and does not entail complex manipulation systems. However, the interaction of acoustics with microstructure geometry is poorly understood to date, and its study is necessary for developing next-generation acoustically powered microrobots. We present here a mass-manufactured acoustically driven helical microrobot capable of locomotion using a fin-like double-helix microstructure. This microrobot responds to sound stimuli and mimics the spiral motion of natural microswimmers such as spirochetes. The asymmetric double helix interacts with the incident acoustic field, inducing a propulsion torque that causes the microrobot to rotate around its long axis. Moreover, our microrobot has the unique feature of its directionality being switchable by simply tuning the acoustic frequency. We demonstrate this locomotion in 2D and 3D artificial vasculatures using a single sound source. Since ultrasound is widely used as an imaging modality in clinical settings, our robotic system can integrate seamlessly into practice; thus, our findings could contribute to the development of next-generation smart microrobots.

**One-Sentence Summary:** We present an acoustically driven helical microrobot capable of corkscrew-like locomotion using a double-helix microstructure.

## INTRODUCTION

Corkscrew motions are ubiquitous among nature’s microscopic swimmers, having evolved in various strains of bacteria, spermatozoa, and other microswimmers as a means of overcoming the viscous drag that dominates at the microscale. For example, Spirochete, a pathogen that causes diseases such as Lyme disease and syphilis, can swim in viscous environments like gel, blood, lymph, or connective tissue through rotation of periplasmic rotors, leading to its overall rotation and corresponding translation *(1, 2)*. Biomimicking such a motion in artificial swimmers is extremely challenging but crucial to the development of next-generation microrobots. The pioneering work of Li and Ghosh et al. has demonstrated helical motion using magnetic actuation *(3, 4)*; however, such motion has yet to be successfully implemented with any other actuation method. Several other pioneering physical and chemical microscale propulsion strategies have been studied, including biohybrids *(5, 6)*, chemical reactions *(7, 8)*, optics *(9)*, enzymes *(10-13)*, electric fields *(14)*, magnetics *(15-19)*, and acoustics *(20-25)*. However, poor biocompatibility, low speed and force, and poor navigation capabilities limit the potential of existing approaches, particularly for medical applications. For example, microrobots that use chemical fuels and electric fields for propulsion may not be entirely biocompatible, as these mechanisms could damage biological tissue. Recently, a promising new possibility has emerged in which enzymes are used to catalyze a reaction resulting in propulsion. However, so far, no controlled manipulation of enzyme-powered microrobots has been demonstrated. Another sophisticated strategy for microrobot propulsion is light activation of photoactive liquid-crystal elastomers; although these can be made biocompatible, light penetrates only to a depth of a few millimeters in human skin, and may be absorbed or scattered by surrounding tissue, making *in vivo* use of this system difficult. Acoustically and magnetically powered micro-/nanorobots are well suited for surgical operations, as both acoustic waves and magnetic fields can penetrate deep into the tissue, are not affected by the opaque nature of animal or human bodies, and generate a broad range of forces that can result in strong propulsive forces *(5, 6, 15-17, 26-29)*. Magnetic microrobots are well-known for their precise navigation capabilities, but require a complex multistep microfabrication process, including doping with magnetic materials; moreover, magnetic manipulation systems are associated with bulky and expensive apparatus. Furthermore, there are strong limitations to the rate of change of the magnetic flux to avoid harmful electromagnetic induction in a medical application.

However, acoustically actuated wireless microrobots, while still in the very early stages of development, are becoming increasingly attractive, as acoustic control can generate large propulsive forces, penetrate deep into the tissue, not being affected by the opaque nature of animal or human bodies, requires a simple microrobot design, and does not entail complex manipulation systems. In the design of ultrasound-based microrobots, it is important to consider the interaction between acoustic waves and (i) material composition and (ii) the geometry of microstructures. For example, trapped microbubbles in polymeric cavities cause intense scattering, bringing about strong propulsive forces when activated by ultrasound. Different sizes and arrangements of microbubbles lead to different motions: rotational and directional. Although acoustically activated microbubbles can generate large propulsive force, their instability limits their practical application *in vivo*. Furthermore, advanced hydrophobic treatments are required to maintain microbubbles within sub-micrometer cavities. To overcome this challenge, researchers have developed bubble-free, acoustically activated micro- and nanorobots. For example, propulsion has been demonstrated through the acoustic oscillation of appendages such as artificial flagella and ciliary bands *(24, 25)*. Wei et al. demonstrated propulsion using bimetallic rods at the pressure nodes of a standing acoustic wavefield *(30)*. This system, however, is intrinsically dependent on the boundaries of a resonating acoustic chamber, which limits its use *in vivo*, and many of its mechanisms remain unknown. Nonetheless, most acoustic systems of propulsion have not been shown to propel in 3D vasculature environments, which is crucial for *in vivo* navigation. Recently, Ren et al. *(31)* and Agakhani et al. *(32)* have demonstrated 3D manipulation of bubble-activated acoustic microrobots; however, these microrobots require gas-filled microbubbles, can only be propelled unidirectionally when they are exposed to ultrasound, and are not able to steer without the application of an external magnetic field. Furthermore, they are difficult to manipulate in vasculature when the microrobots’ length exceeds the diameter of the vessel; thus, a microrobot design is needed that can induce bidirectional motion without physical orientation, which is necessary to avoid occluding the vessels. To introduce novel acoustic propulsion mechanisms with new functionalities, we need to investigate the ways in which sound interacts with the microarchitectures of various geometrical configurations. At present, the interaction between acoustic wavefields and complex geometrical microstructures, particularly in the context of propulsion, has almost not at all been studied.

In this article, inspired by the helical geometry of *spirochete* bacteria, we present a mass-manufacturable acoustically driven helical microrobot (or micro-propeller) that is capable of locomotion using a fin-like double-helix vane. This microrobot responds to external sound stimuli and mimics the spiral motion used by spirochetes and spirilla, which are natural microswimmers. The asymmetrical double helix shape serves to interact with the incident acoustic field, inducing a propulsion torque that causes the microrobot to rotate around its long axis. Moreover, our microrobot has the unique feature of a propulsion direction that is switchable simply via tuning the acoustic frequency. We demonstrate this locomotion in artificial arbitrarily shaped 2D and 3D vasculatures using a single sound source. Because ultrasound is already in widespread use as an imaging modality in clinical settings, our robotic system can be seamlessly integrated into its practices; thus, our findings could contribute significantly to the development of next-generation smart microrobots for non-invasive surgical procedures.

## RESULTS

### Particle design

Inspired by the locomotion of helical natural microswimmers such as *Spirochete* bacteria (Fig. 1A), we have designed a wireless microrobot consisting of a cylindrical core and a double-helical vane that coils around the core (see Fig. 1). The double-helical vane can be seen as a superposition of two curved vanes that spiral along the surface of the cylindrical core as helicoids. When this microrobot is surrounded by a fluid like water and exposed to an acoustic field, it is propelled forward by a corkscrew-like motion, where the particle rotates around the axis of its cylindrical core and simultaneously translates along this axis. Our microrobots can be mass-manufactured by a high-resolution two-photon lithography 3D printing technique. The microrobots that we have manufactured for the present study have length 350 μm and diameter σ =100 µm and are made from a polymeric material (see Materials and Methods).

**Fig. 1.**
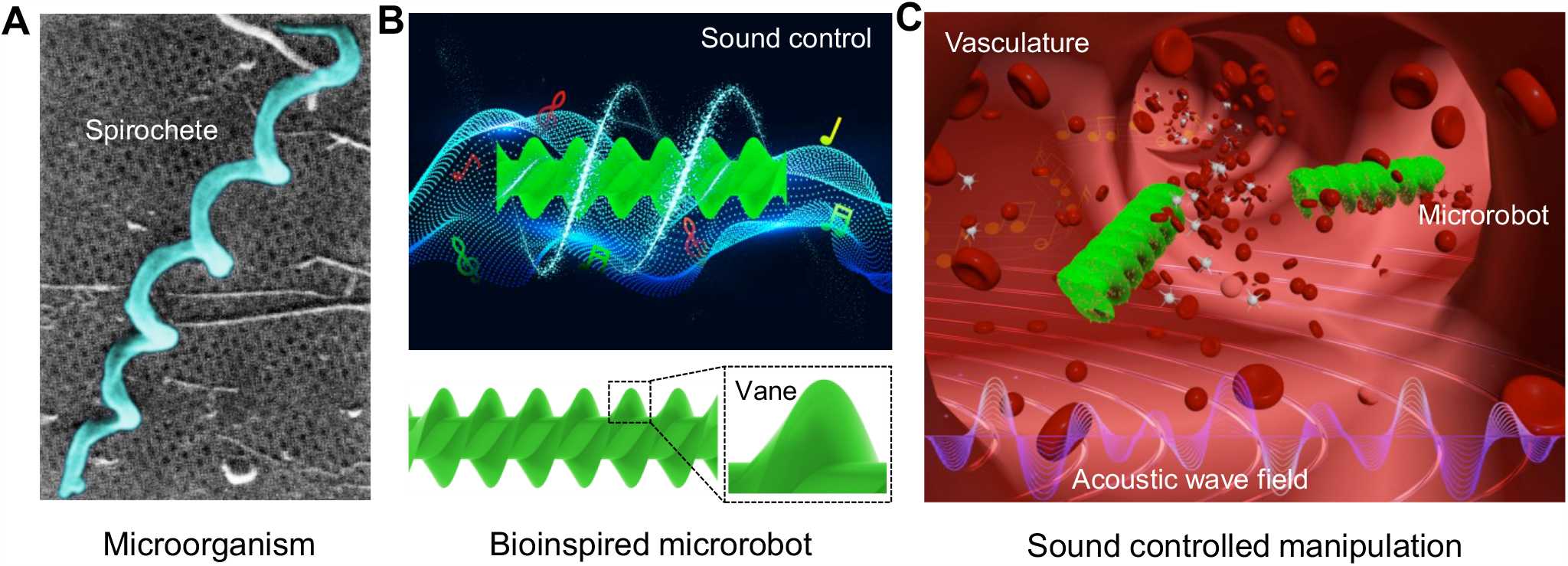
Spirochete-inspired propulsion based on acoustic actuation. (**A**) Micrograph showing the spiral morphology of *Spirochete (33)* bacteria, which execute translation motion through rotation in viscosity-dominated fluids — reprinted with permission from Ref. 33. Copyright ^a^ 2018 American Chemical Society. (**B**) An illustration of a wireless bioinspired robot that propels in response to an external sound field. (**C**) A concept schematic illustrates the use of such microrobots for non-invasive surgery in vasculatures.

### Propulsion mechanism

To determine the forces and torques exerted on these microrobots and thus to characterize their propulsion mechanism, we performed acoustofluidic computer simulations. Starting from the compressible Navier-Stokes equations

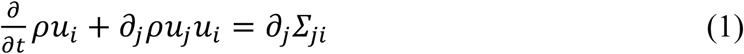

with the momentum-stress tensor

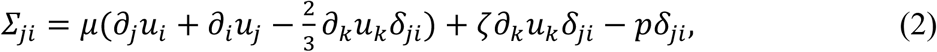

where *t* denotes time, *∂*_*i*_ denotes a partial derivative with respect to the *i*th spatial coordinate, *u*_*i*_, *ρ*, and *p* are the fluid’s velocity, mass density, and pressure fields, *μ* and *ζ* are its constant shear- and bulk viscosities, *g* represents additional source terms, and *δ* is the Kronecker-Delta symbol. By using the continuity equation as an additional condition, the total force *F*_*i*_ and torque *T*_*i*_ on the particle can be expressed as an integral over the particle volume *Ω* as

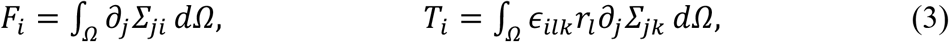

where *ϵ* is the Levi-Civita symbol and *r*_*i*_ is the location of the volume element *dΩ* relative to the microrobot’s center. We do not consider wave propagation through the material of the particle. Therefore, Stokes’ theorem can be used to rewrite *F*_*i*_ as an integral over the particle surface *∂Ω* as

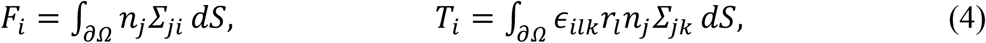

where *n*_*i*_ is the normal vector of the surface element *d∂Ω* = *n*_*i*_*dS (34)* with the surface differential *dS*. Since the structure of the microrobot has no translational invariance, a full 3D simulation is needed to investigate the microrobot’s propulsion mechanism. Furthermore, a large domain in space and time (multiple hundred wave periods to reach a sufficiently converged solution *(35)*) needs to be covered with high resolution, especially around the particle, and with a small temporal step size *(36)*.

Using the full compressible Navier-Stokes equations as the governing equations, such a simulation would be computationally extremely expensive. To avoid this, the Nyborg expansion *Σ* − *Σ*_0_ = *Σ*_1_ + *Σ*_2_ + ⋯ with

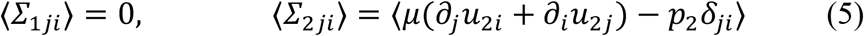

is used to finally arrive at the simplified expressions for the radiation force and torque *(34, 37-41)*

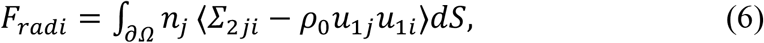

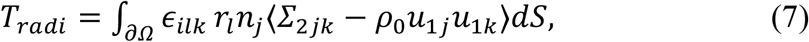

where ⟨⟩ denotes a time average over one wave period. The subscripts 0, 1, and 2 denote different orders in the Nyborg expansion and the time-averaged momentum-flux tensor ⟨*ρ*_0_*u*_1*j*_*u*_1*i*_⟩ serves as an approximation for the streaming force *(42)*. A more in-depth view of this derivation can be found in **Supplementary Note 1**. As we now only consider the locally time-averaged solution of a linearized problem, the system can be solved more efficiently. To solve the system of equations numerically, the COMSOL Multiphysics solver was used. See **Methods** for the parameter values chosen in our simulations.

The results of the 3D simulations show the propulsion under sound exposure (see Movie S1). Results for the propulsion force and torque exerted on a particle are shown in Fig. 2 (F and G). As can be seen, the propulsion force and torque are primarily generated by the asymmetrical vanes, with minimal contribution from the end faces and cylindrical core of the microrobot. Due to the asymmetry of the helical fin-like vanes, the acoustic field around the particle interacts with the particle surface in a way that invokes a torque around the particle’s main axis, as illustrated in Fig. 2D. The subsequent non-reciprocal corkscrew-like motion around this axis (depicted in Fig. 2E) can be observed in all experiments alongside the translational motion. Thus, the simulation results show that the rotation of the particle is not a side effect of translational propulsion in combination with a helical particle shape. Instead, there is a significant propulsion torque that leads to the particle’s rotation directly and, in combination with the helical particle shape, can contribute to the particle’s translational propulsion. In a low Reynolds number fluid (the Reynolds number *Re* = *ρuσ*/*μ*, with the characteristic length scale *σ* = 100 *μ*m, is *Re* < 0.03 ≪ 1), the second-order steady flow (also known as acoustic streaming, see Supplementary Information for details) may play a role in propulsion. However, we demonstrate that the acoustic streaming is very weak and can be ignored in our case where the local flow is caused by the locomotion of the microrobot (see fig. S2 and Movie S2). Thus, the acoustic radiation force and rotation of the microrobot are likely to dominate the propulsion process. In the following paragraphs, we address the connection between the translational and rotational motion of the microrobot in more detail.

**Fig. 2.**
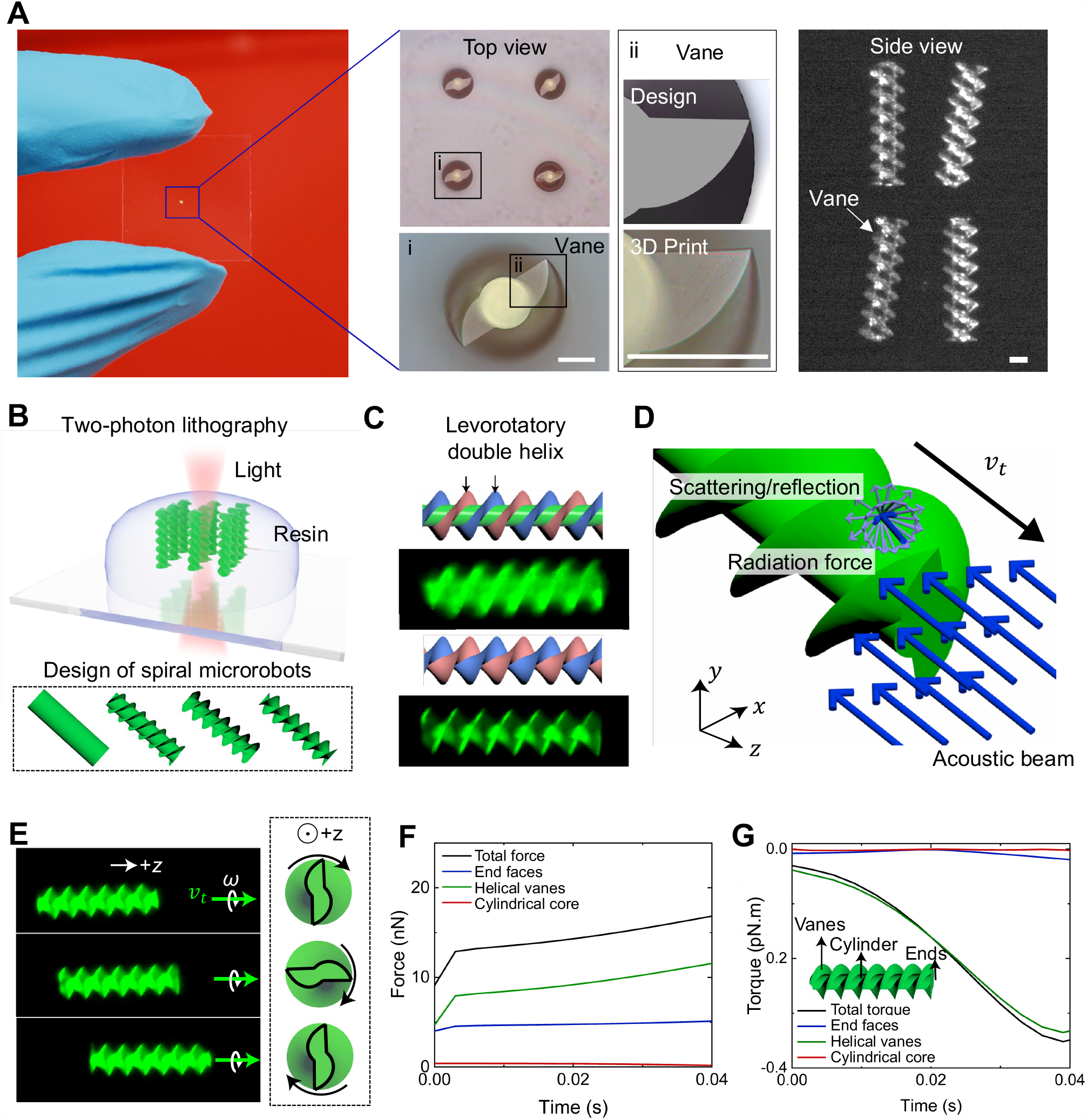
Design, fabrication, and concept of the acoustically actuated helical microrobot. (**A**) Micrograph illustrating the fabricated microrobots on a thin glass slide. The inset shows a top view (left panel) and side view (right panel) of the fabricated microrobots. Scale bar, 40 μm. (**B**) The acoustic helical microrobots are mass-manufactured using the two-photon lithography technique. The bottom inset shows different designs of the helical microrobots. (**C**) Schematic and fluorescent micrographs illustrating the helical vane structures on the microrobot. The pink and blue colors represent the two respective levorotatory double-helix vanes developed on the microrobot. (**D**) Schematic illustrating an asymmetric acoustic force distribution on the surface of the microrobot. (**E**) Image sequence demonstrating the acoustic microrobots’ translational and rotational motion in response to an acoustic stimulus. The insets show the clockwise rotation of the microrobot, which is indicated by the orientation of the vane geometry. (**F** and **G**) Plots of the acoustic radiation forces (**F**) and torques (**G**) generated by the end faces, helical vanes, and cylindrical core components of the microrobots based on COMSOL Multiphysics simulations for sound frequency 13.5 kHz and pressure amplitude 60 kPa.

The fact that, in all experiments, fast translational motion could be observed together with rotation around the particle’s main axis indicates strongly that the geometrical shape of the microrobot results in a coupling between translational and angular propulsion. Intuitively, this mechanism is caused by the hydrodynamic resistance of the microrobot’s helical fin structure. To test this assumption, we compared the observed translation velocity of a microrobot with the translation velocity of a particle with the same shape whose translation results from a purely rotational propulsion through the translational-rotational coupling in the particle’s hydrodynamic resistance matrix.

In a liquid at a low Reynolds number, the hydrodynamic resistance matrix can be used to transform between a particle’s velocity and the force or torque acting on the particle that causes this velocity:

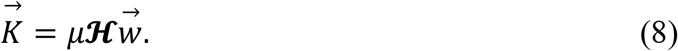

Here, 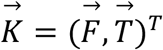 is the force-torque vector that acts on the particle, ***ℋ*** is its shape-dependent 6×6-dimensional hydrodynamic resistance matrix, and 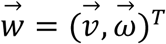 is the particle’s translational-angular velocity vector with the translational velocity 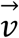 and the angular velocity 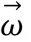. To determine ***ℋ***, we used the software *HydResMat (43, 44)* that solves the Stokes equation using the finite element method.

Since the main rotation action of the particles in the experiments is observed around a particle’s main axis, it can be concluded that the main torque acting on the particle acts along this axis. Assuming a force-torque vector 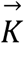, where all vector components except for *T*_*z*_, which is the torque component parallel to the main axis, are zero, we can find a value for *T*_*z*_ such that the corresponding angular velocity *ω*_*z*_ matches that observed in the experiments. In this way, an approximation is gained for both the propulsion torque of the particle and the component of the translation velocity resulting purely from the particle’s rotational motion.

Using the hydrodynamic resistance of the particle, the maximum rotation speed of *ω*_*z*_ ≈ 1.66 Hz observed in the experiments results in a translational velocity of *v*_*rtz*_ ≈ 126 *μ*m/s which only slightly differs from the translational velocity of *v*_*tz*_ ≈ 111.9 *μ*m/s that we observed in the experiments. The corresponding torque along the main axis of the particle is *T*_*z*_ ≈ 618 nN*μ*m. Repeating this comparison between the theoretically predicted rotation-induced translation velocity and the translation velocity observed in experiments for particles with slightly differing geometrical properties allows for a quantitative assertion of how significant the contribution of the particles’ rotation to its translational motion is. The results of these comparisons are plotted in Fig. 4 (B and D) and show a strong coherency between the rotation-induced translational velocity *v*_*rt*_ introduced purely through hydrodynamic translational-rotational coupling and the total translational velocity *v*_*t*_ observed in the experiments. This indicates that the rotation about the main axis of the particle is a major contributor to its translational motion. The small discrepancy between *v*_*rt*_ and *v*_*t*_ can be explained in part by acoustic radiation applying a driving force to the particle directly and by drag between the particle and the channel walls in the experiments, which has not been taken into account for the calculation of ***ℋ***. The evidently strong coupling between acoustically induced rotation and translation is a unique feature to our particle design.

### 2D manipulation

We performed 2D manipulation of our microrobots in a rectangular acoustic chamber filled with water and in a microchannel with a circular cross-section filled with an ethanol solution. The acoustic chamber and microchannel were made of PDMS and, together with a piezoelectric transducer, bonded onto a glass slide. The piezoelectric transducer, which generates the sound field driving the microrobots, is regulated by an external function generator connected to an amplifier. The entire setup was then mounted on a microscope, and high-speed and high-sensitivity cameras were used to record the experiments to analyze the mechanism behind the propulsion (see details in Materials and Methods).

For the microfluidic channel, we chose a circular cross-section of 500 μm in diameter, a size that mimics the diameter of the average human blood vessels. In response to an acoustic field, the microrobot advances unidirectionally along the channel at a uniform speed while rotating about its *z*-axis. Fig. 3A shows the left-to-right locomotion of the microrobot at 100 μm/s accompanied by clockwise rotation at 1.66 *s*^−1^ when the microrobot is exposed to an acoustic field with frequency *f*_1_ = 13.5 kHz and peak-to-peak voltage 20 V_pp_. As we modulate the driving frequency of the acoustic signal, the microrobot shows a counterclockwise rotation and translational motion in the opposite direction. Fig. 3B illustrates the right-to-left motion of the microrobot at ≈70 μm/s and rotation at 0.9 *s*^−1^ for *f*_2_ = 18.6 kHz and 60 V_pp_. Fig. 3C shows this bidirectional propulsion along a channel. Upon activation at *f*_1_, the microrobot is translated along the microchannel at ≈100 μm/s. After 13.2 s, we turned off the acoustic signal, causing the microrobot to roll back to the middle of the microchannel due to gravity. At 22.5 s, when the transducer was activated again at *f*_2_, the microrobot was propelled in the opposite direction at ≈70 μm/s. This ability to alter the directionality of the microrobot by almost 180° by simply switching the activation frequency of the acoustic signal is a unique feature that can potentially avoid occluding vessels when acoustically actuated microrobots are applied in medicine.

**Fig. 3.**
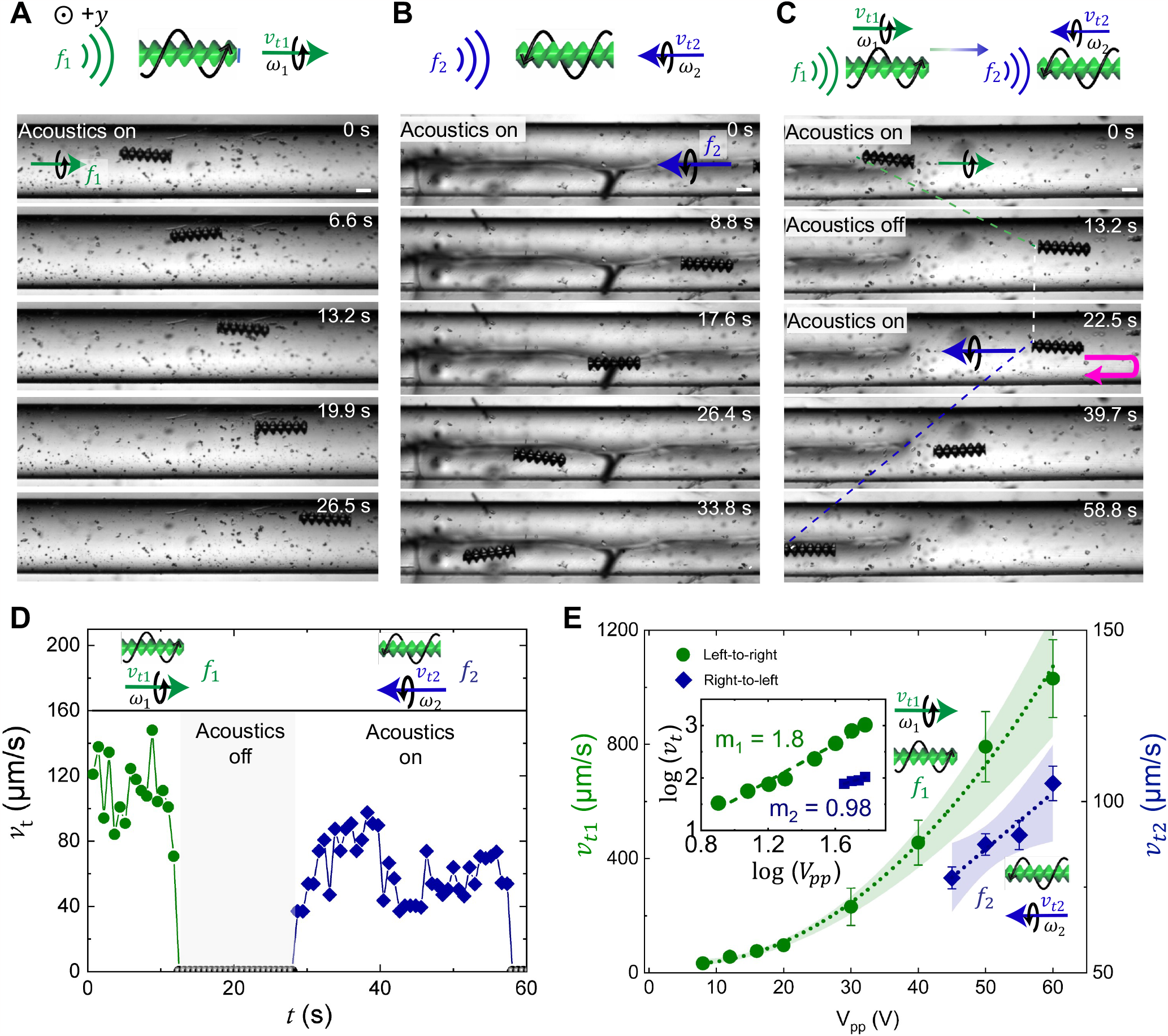
Translational motion of the microrobot in a circular microchannel. (**A**) The microrobot exhibits left-to-right locomotion in response to an external acoustic wave at *f*_1_ = 13.5 kHz and 20 V_pp_. (**B**) The microrobot exhibits right-to-left locomotion for *f*_2_ = 18.6 kHz and 60 V_pp_. (**C**) A microrobot illustrates its bidirectionality when switching from *f*_1_ = 13.5 kHz and 20 V_pp_ to *f*_2_ = 18.6 kHz and 60 V_pp_. (**D**) The plot illustrates the speed profile of the microrobot during its bidirectional trajectory. (**E**) The microrobot’s left-to-right velocity *v*_t1_ and right-to-left velocity ***v***_**t2**_ versus the acoustic driving voltage V_pp_ indicate that the microrobot propels at a speed nearly proportional to 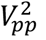. The inset shows the corresponding log-log plot. Note that the scaling of *v*_t2_ was difficult to predict due to a lack of data points at higher voltages, since the maximum voltage is restricted by our amplifier at 60 V_pp_. Scale bar, 100 μm. The direction of gravity is antiparallel to the *y* direction (indicated by ⨀), i.e., perpendicular to the figure plane. Each data point represents the average velocity of at least 3 microrobots. The error bar represents the standard deviation.

The unidirectional and bidirectional trajectories of the microrobot are consistent both in the rectangular chamber (fig. S3 and Movie S3) and in the microchannel (Movie S4), indicating that its behavior is determined by the acoustic field and not significantly influenced by the channel wall. The microrobot executed cork-screw motion, i.e., rotation combined with translation in a straight trajectory in both open and closed channels. However, the excitation frequencies varied by 11% between the rectangular chamber and open channel, which could be attributed to differences between the water and ethanol solutions in which the microrobots were submerged. Interestingly, we also observed that the left-to-right (i.e., parallel to the sound propagation) propulsion velocity is generally ≈ 40% greater than the right-to-left (i.e., antiparallel to the sound propagation) propulsion velocity, independent of the boundary condition. Likely reasons for this difference are that the acoustic radiation force depends on the frequency and that the propulsion might have a component that is parallel to the propagation direction of the sound in both cases. We further measured the speed of the microrobot as a function of the intensity of the incident acoustic field, controlled by adjusting the voltage applied to the piezoelectric transducer. The propulsion velocity *v*_*t*_ scales quadratically with the applied voltage 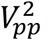 i.e., *v*_*t*_ ∝ 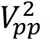, as shown in Fig. 3e. This scaling relationship is expected, since the acoustic pressure amplitude is proportional to *V*_*pp*_ and the acoustic energy density is proportional to the pressure amplitude squared. Numerical and experimental studies have shown that the pressure amplitude depends linearly on applied voltage for small signal power values *(45)*.

The consistent coherency of rotational and translational velocity signifies the fact that the rotation of the microrobot contributes to translation. Approximating the torque experienced by the particle through hydrodynamic resistance, calculated from the hydrodynamic resistance matrix ***ℋ*** using the observed terminal angular velocity, yields a translational velocity of ≈126.0 μm/s purely through particle rotation and hydrodynamic resistance, without taking account of any other effects. This fact also indicates that particle rotation is a main contributor to the overall particle velocity. The difference between the observed translational velocity and the one calculated for rotational-translational coupling through hydrodynamic resistance can in part be explained by the absence of friction forces between the particle and the channel walls in the computational model.

### Structure-dependent behavior of the microrobot

In this section, we investigate the parametric effects of the width and pitch of the helical vane structure on the microrobot’s propulsion. To characterize the width, we introduce as a dimensionless parameter the effective radius ratio (*r*_*d*_ = *r*/*R*), which describes the ratio of the radius *r* of the microrobot’s cylindrical core to the microrobot’s total radius *R*. First, we fabricated a cylindrical microstructure without any helical vanes (i.e., *r*_*d*_ = 1), which also served as a control in that, when subjected to acoustic stimulation, the microstructure undergoes no rotational motion and nearly no translational motion. We then designed microrobots with helical vanes with *r*_*d*_ values of 0.65, 0.43, and 0.22. The design and cartoon schematics of the microrobots are shown in the left panel of Fig. 4A. As *r*_*d*_ decreases, the microrobots’ vanes become wider, resulting in faster rotation rates and higher translation velocities at constant activation parameters of 13.5 kHz and 20 V_pp_, as shown in Fig. 4A. The micrographs shown in Fig. 4A represent the distances travelled by these different microrobots over 10 s of acoustic stimulation; the microrobot with the lowest *r*_*d*_ value (*r*_*d*_ =0.22) travelled the farthest, see also Movie S5. Fig. 4B estimates the translational velocity *v*_*t*_ and rotational velocity *ω* of a microrobot as functions of *r*_*d*_. Lower *r*_*d*_ values result in larger values of *v*_*t*_ and *ω*, which can be attributed to the larger dimensions of the vanes. We further calculated, based on the microrobots’ hydrodynamic resistance matrix (see Propulsion mechanism for details), their translational velocity *v*_*rt*_ that corresponds to their rotational velocity *ω* as a function of *r*_*d*_ (see Fig. 4B). The result for *v*_*rt*_ is very similar to *v*_*t*_, indicating that the microrobots’ rotation is crucial for their propulsion.

**Fig. 4.**
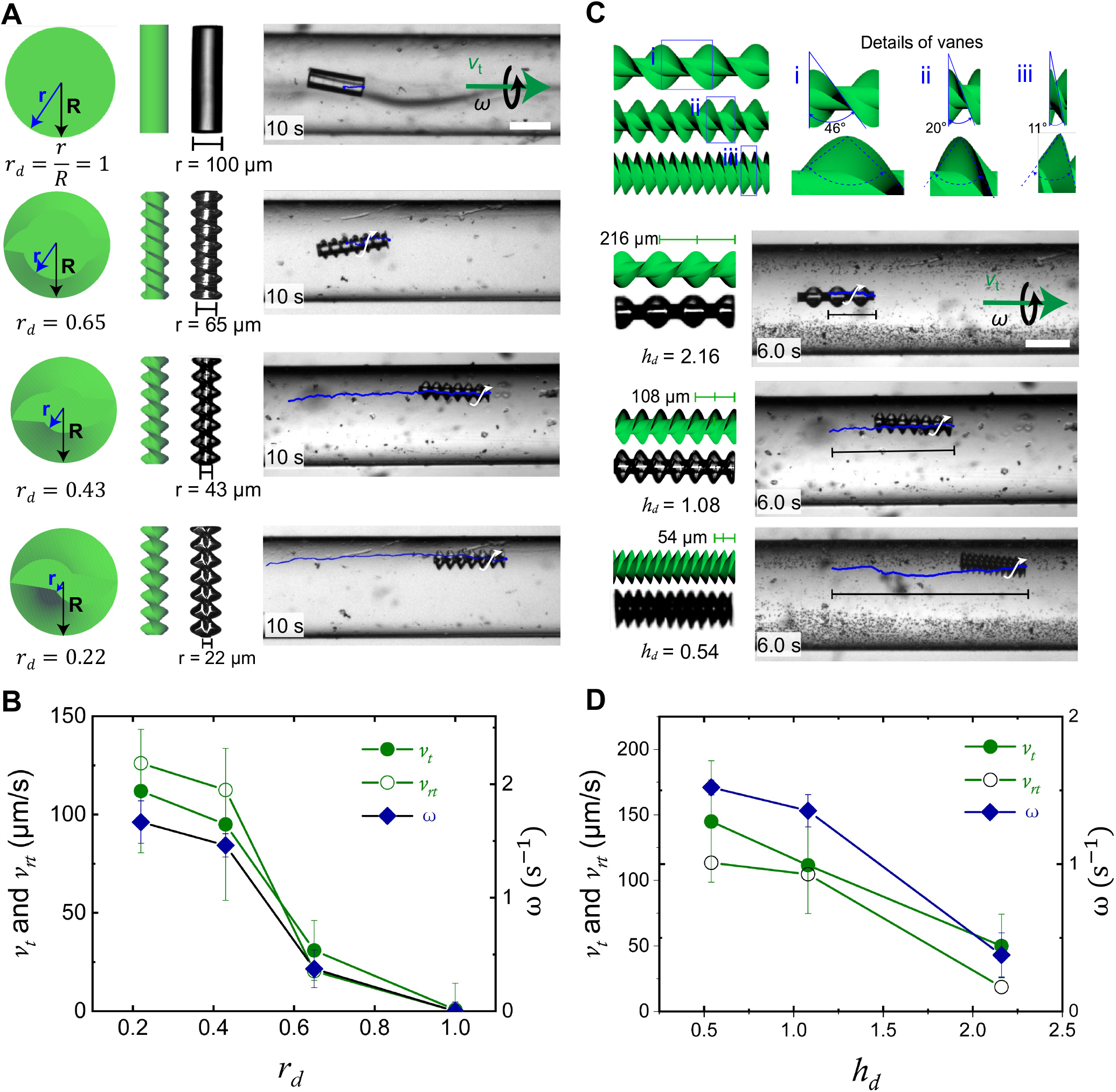
The structure-dependent behavior of the microrobots. (**A**) Parametric study of the effect of the effective diameter ratio (*r*_*d*_) on the translation and rotation velocities of the microrobot. With decreasing *r*_*d*_ values, microrobots exhibit faster translation, which can be attributed to increases in their vane width and surface area. The left panel illustrates the design of microrobots with *r*_*d*_=1.0, 0.65, 0.43, 0.22. The right panel demonstrates the distance travelled for each corresponding *r*_*d*_ value over 10 s at the constant activation parameters of 13.5 kHz and 20 V_pp_; see also Movie S5. Scale bar, 200 µm. (**B**) Plot of a microrobot’s translational velocity *v*_*t*_ (green), angular velocity *ω* (blue), and rotation-induced translational velocity ***v***_**rt**_ (white) as functions of *r*_*d*_. (**C**) Parametric study of the effect of the helical pitch (*h*_*d*_) on the translation and rotation velocities of the microrobot. The left panel illustrates the design of microrobots with *h*_*d*_=2.16, 1.08, 0.54. The right panel demonstrates the distance travelled for each corresponding *h*_*d*_ value over 6 s at 13.5 kHz and 20 V_pp_; see also Movie S6. Scale bar, 200 µm. (**D**) Plot of the microrobot’s translational velocity ***v***_**t**_ (green), angular velocity ***ω*** (blue), and rotation-induced translational velocity ***v***_**rt**_ (white) as functions of *h*_*d*_.

Next, we characterized the effect of the reduced helical pitch *h*_*d*_ = *P*/(2*R)* on the microrobots’ propulsion, where *P* is the helical pitch. We fabricated microrobots that had *h*_*d*_ values of 2.16, 1.08, and 0.54. As *h*_*d*_ decreases, the number of helical turns on the microrobot’s surface increases, as demonstrated in Fig. 4C (see also Movie S6). Microrobots with lower *h*_*d*_ values exhibited faster angular and translational propulsion velocities when exposed to acoustic stimulation; a microrobot with *h*_*d*_ = 0.54 was propelled the most rapidly at ≈150 μm/s. Fig. 4D indicates the translational velocity *v*_*t*_ and the rotational velocity *ω* attained by the microrobots as functions of *h*_*d*_. As *h*_*d*_ decreases, both *v*_*t*_ and *ω* increase, which might be attributed to the larger number of helical turns of the vanes. We further calculated the microrobots’ translational velocity *v*_*rt*_ from *ω* as a function of *h*_*d*_, which supported the importance of the microrobots’ rotation for their translational propulsion.

### 3D manipulation

Microrobots that can deliver drugs directly to lesions through the vasculature are the next generation of drug carriers. To execute this task, microrobots should be able to navigate in 3D. In this section, we studied the microrobots’ propulsion and navigation behavior in response to acoustic stimulation in arbitrary 3D vasculatures. We created PDMS-based microchannels with various angles (0°, 15°, 25°, 45°, 60°, and 75°), defined as the angle of inclination of the microchannel to the horizontal plane, as shown in Fig. 5A (see also fig. S4). These PDMS-based microchannels are fabricated by molding electric wires into PDMS, which are removed after curing. Fig. 5B shows a photograph of a PDMS-based microchannel. The concept of bidirectional motion when the excitation frequency is switched from *f*_down_ (downwards) to *f*_up_ (upwards) is shown in Fig. 5C. Fig. 5D shows superimposed images of microrobots at different times during downward motion in an acoustic field at 15°, 25°, 45°, 60°, and 75° inclination. The microrobots executed rotation along their long axis and travelled at an almost uniform velocity, see also Movie 7. When we increased the slope from 45° to 60°, no noticeable change in behavior was observed. As the microrobot is propelled through channels with larger slopes 45°, 60°, and 75°, it becomes blurry or out-of-focus, since we kept the focus of the microscope constant. This effect is more pronounced with 75° slope at the right of the bottom panel in Fig. 5D.

**Fig. 5.**
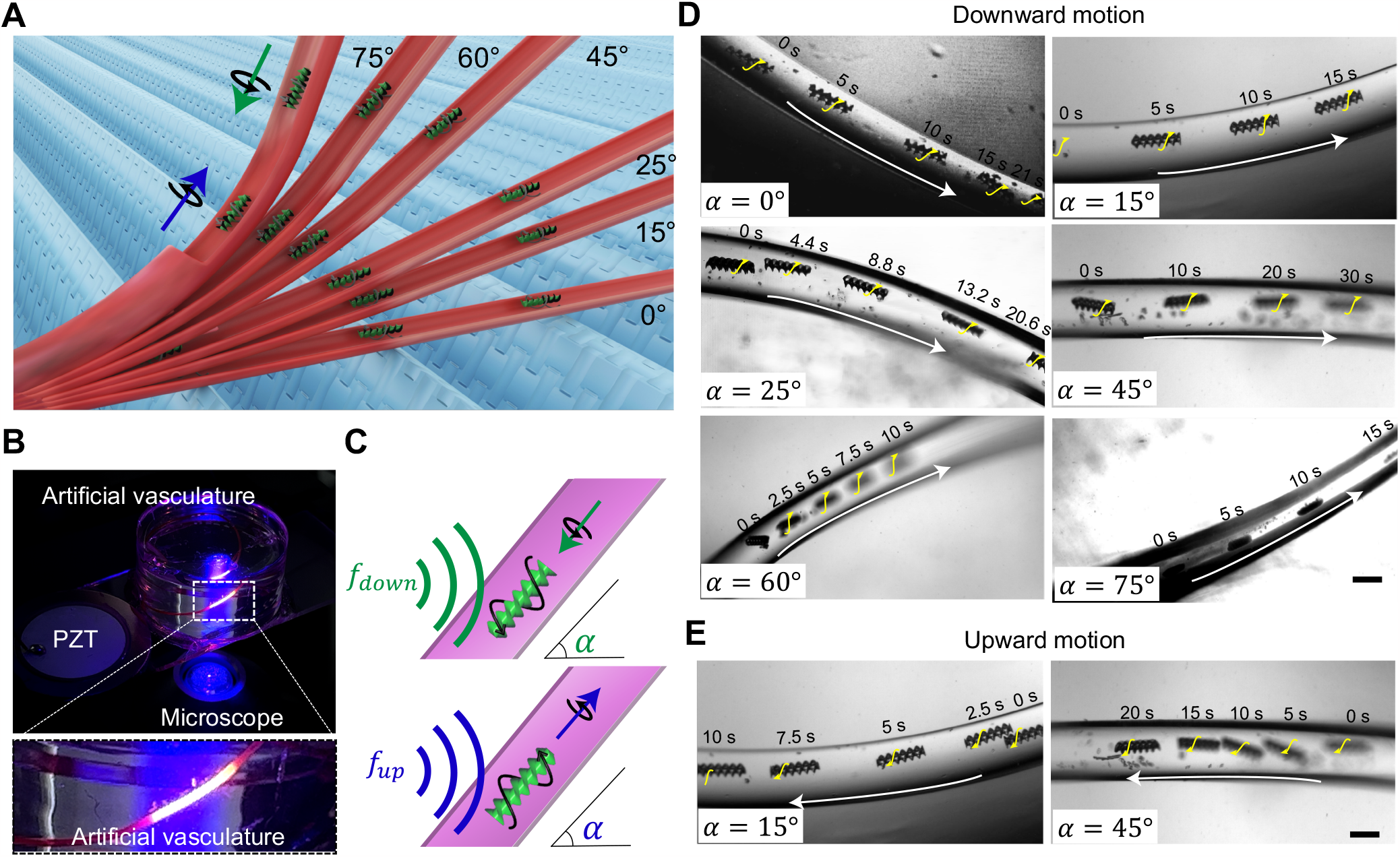
Manipulation of the microrobots in a 3D artificial vasculature. (**A**) A schematic illustrates the manipulation of acoustic helical microrobots in channels with different angles, see also fig. S4. (**B**) The photograph depicts a 3D PDMS-based artificial vasculature mounted on a glass slide adjacent to a piezo transducer. The inset shows the fluorescently labelled channel with a circular cross section. (**C**) Concept of bidirectional motion when the excitation frequency is switched from *f*_down_ (downwards) to *f*_up_ (upwards). (**D**) Superimposed images showing downwards motion (*f*_down_=11.7 to 13.7 kHz) of microrobots in 3D channels with angles of 0°, 15°, 25°, 45°, 60°, and 75°. (**E**) Examples of superimposed images showing an upward propulsion (frequency *f*_up_ =14.5 to 15.1 kHz) of microrobots in 3D channels with angles of 15° and 45°. See also Movie S7 and S8. Scale bars, 250 μm.

In addition to its downward motion, the developed microrobots also exhibited controlled upward motion in a microchannel, as shown in Fig. 5E. The microrobot changed the direction of its rotational and translational motion as the stimulation frequency (*f*_up_=14.5−15.1 kHz) was switched in microchannels at inclinations 15° and 45°, indicating that bidirectionality of the microrobots is preserved in 3D vasculatures (Movie S8). The time stamps in Fig. 5E indicate that the microrobot in a 15° slope vasculature propels much faster than a 45° slope vasculature, thus indicating that the microrobot is being propelled in 3D against gravity. The 3D manipulation and bidirectionality of acoustically actuated microrobots pose a bottleneck in many situations; our observations indicate that our microrobots offer a promising solution for this issue.

## DISCUSSION

This study presents a biomimetic, acoustically controlled, spiral-shaped microrobot inspired by *Spirochete* bacteria. The microrobot responds to ultrasound stimuli from outside the body and mimics the spiral motion of natural microswimmers to execute propulsion at low Reynolds numbers. They exhibit simultaneous rotation and translation motion, enabled by the interaction between acoustics and the double-helix microstructure. The microrobots have exhibited bi-directional capabilities, meaning that they can propel even antiparallel to the propagation direction of the sound wave, and have a three-dimensional range of motion using a single acoustic piezo transducer. Magnetic microrobots with helical structures have been reported, but the underlying mechanics of acoustically activated microrobots and the related propulsion concept are fundamentally different *(4, 18, 46-49)*. Magnetic microrobots are often restricted to a step-out frequency (typically in the Hertz regime), where the magnetic torque applied is unable to keep the microrobot synchronized with the external magnetic field. This problem cannot be solved by increasing the magnetic flux density, since electromagnetic induction must be kept at a harmless level in a medical application. In this context, acoustically actuated microrobots can attain faster propulsion because their rotation velocities scale quadratically with input voltage and can reach larger values already for harmless ultrasound intensities. Future research will focus on faster rotation-induced translational motion based on optimizing the microrobot’s geometry, size, and materials.

The microrobots’ translational-rotational coupling which transforms a strong rotational propulsion into a fast translation is a unique feature compared to previous realizations of acoustically actuated microparticles. These microrobots can be designed to rotate and thus translate more quickly by incorporating microbubbles on one side of the vane. When these microbubbles are acoustically activated at resonance, they can produce a stronger propulsion as is needed to allow the microrobots to propel through a gel-like medium.

Future research will examine the microrobots’ steering capabilities. At present, our microrobots can propel in artificial vasculature with bidirectional capability and perform three-dimensional manipulation. When microrobots perform tasks *in vivo*, they may encounter a number of challenges, including head-to-tail inversion in tiny vessels. When the particles are long compared to the vessel diameter, they can cause an occlusion by reorientation or be not able to invert their orientation and thus propulsion direction at all. Since our microrobots can move bidirectionally without head-to-tail inversion by tuning the excitation frequency, we can prevent these problems. We also look forward to having our microrobots navigate through smaller blood vessels and capillaries. Microstructures with sub-micron helical characteristics can be easily fabricated with 2D photon lithography. When scaling down the microrobots, the underlying propulsion mechanism will require further study. In the current design of the microrobots, acoustic streaming has little effect on propulsion. However, as we scale the microrobots down, the dominance of the acoustic radiation and streaming forces will change. Future work will examine how the microrobots’ size affects their propulsion.

The concept of the rotational and translational motion of microrobots can be further used in the development of wireless stents. For instance, we can design a stent with a similar helix structure and use acoustics to implant it into the lesion area through acoustic manipulation. Future microrobots can be more functional for more flexible and customized biomedical applications. For example, drug-containing microbubbles can be equipped with our microrobots in the process of drug delivery by using transducers with different excitation frequencies to prevent interference between their propulsion and drug-delivery functions. Also, as our microrobot is manipulated by acoustics, integrating it with clinical ultrasound may be possible for simultaneous manipulation and monitoring of its motion. The developed microrobots can also trap and transport micro- to millimeter-sized objects, such as cells, organoids, tissue constructs, and model organisms like *Caenorhabditis elegans* and zebrafish embryos to desired locations using the scattered sound fields of the microrobots. Furthermore, the microrobots can be built with hollow cavities to trap and transport sperm cells for *in vitro* fertilization.

## MATERIALS AND METHODS

### Fabrication of helical microrobots

The acoustics-driven helical microrobots were directly mass-fabricated according to a standard procedure by Nanoscribe (Nanoscribe Photonic Professional GT, Nanoscribe GmbH) based on two-photon 3D laser lithography (24). This technique allowed the direct and precise printing of complex 3D micro/nano-structures at an enhanced resolution. A drop of photoresist (IP-Dip, Nanoscribe GmbH) was placed on an IPA-rinsed (isopropyl alcohol) standard glass slide (25 mm × 25 mm × 1 mm), which was precisely inserted into a microscope system for laser writing through a 63× objective. The printing system could expose and cure the photoresist programmatically until it was completed, as shown in Fig. 2B. After writing, the glass slide with microrobots on was carefully rinsed three times using acetone and IPA for 10 min each. The printed microrobots were transferred into a microchannel for experiments using a 140 μm-diameter glass fiber.

### Manipulation of the microrobot

Experiments of 2D and 3D manipulation were carried out in a rectangular channel (20 mm × 20 mm) or circular microchannel in PDMS (Dow Corning). The 2D manipulation meant the microrobot was maneuvered horizontally by modulating the frequency in a horizontally straight channel. We fabricated the rectangular channel and circular microchannel by using a piece of glass and a wire as a mold (500 μm in diameter) in a PDMS block, respectively. PDMS curing occurred in a 3D-printing mold in a heater at 80°C for 2 h. A glass slide (75 mm × 25 mm × 1 mm) coupled with such PDMS and a transducer (Steiner & Martins, Inc., USA) was mounted on an inverted microscope (Carl Zeiss, Germany) with various objectives (2.5x, 5x, 10x, 20x) for imaging, as shown in fig. S1. The manipulation system, mainly consisting of a function generator (0 – 10 MHz, 0 – 20 V_pp_, Tektronix, Inc., USA) and an amplifier (0 – 60 V of output, 15-time magnification, Digitum-Elektronik, Germany), generated a parameter-adjustable acoustic wave (wave types, voltage, and frequency). A continuous square wave was applied. The imaging system, including a microscope and a camera, was used to record the time-lapse images for further analysis. A normal camera (Photometrics, USA) and a high-speed camera (Kron Technologies Inc., Canada) were alternatives. A microrobot was transferred into the microchannel by a fiber, and a solution of Isopropyl alcohol was injected as solvent.

With respect to 3D manipulation, experiments were carried out in a 3D artificial vasculature or microchannel in a PDMS block. Similarly, the 3D microchannels were fabricated in PDMS using a curled wire of diameter 500 μm to make the microchannels more complicated. Different types of microchannels, including angles of 0°, 15°, 25°, 45°, 60°, and 75°, were fabricated as shown in Fig. 5B and fig. S4. Then, the transducer and PDMS were coupled on a rinsed glass slide carefully, and a microrobot was transferred into the 3D microchannel for subsequent experiments. Isopropyl alcohol was used as the solvent.

### Acoustofluidic simulations

COMSOL Multiphysics (Version 5.6) was used to solve the relevant acoustofluidic equations in a 3D environment in this study. A CAD file of a microrobot (350 μm in length and 100 μm in diameter) was imported and a microchannel of diameter 500 μm was constructed as a flow domain. The modules Thermoviscous acoustics, Laminar flow, and Solid mechanics were coupled to simulate the acoustic field, fluid field, and stress field in the particle. Applying suitable boundary conditions, the acoustic pressure field was calculated, based on which the acoustic streaming and second-order acoustic radiation force were determined using the perturbation method. Using Eqs. (6) and (7), we then calculated the acoustic propulsion force and torque.

## SUPPLEMENTARY MATERIALS

Movie S1: Simulations of the microrobot.

Movie S2: Propulsion of the microrobot in 2D channel.

Movie S3: Bidirectional propulsion of the microrobot in a rectangular chamber.

Movie S4: 2D bidirectional propulsion of the microrobot in the artificial vasculature (circular-cross-section microfluidic channels).

Movie S5: Effects of the effective radius ratio r_*d*_ on the translation and rotation velocities of the microrobot in the microchannel.

Movie S6: Effects of the reduced helical pitch *h*_*d*_ = *P*/(2*R)* on the microrobot’s propulsion.

Movie S7: 3D manipulation of the microrobot in the microchannels with different inclinations (0°, 15°, 25°, 45°, 60°, and 75°).

Movie S8: 3D upward and downward manipulation of the microrobot in microchannels inclined at 15°.

Note 1. Numerical simulations.

References (50-52)

## Acknowledgments

We thank Mahmoud Medany for the assistance of experiments.

## Author contributions

D.A. conceived and supervised the project. Y.D., Z.Z., and D.A. contributed to the experimental design. Y.D. and Z.Z. performed the experiments. Y.D., Z.Z., and A.P. performed the computer simulations. Y.D., A.P., and R.W. contributed to the theoretical understanding. Y.D., Z.Z., A.P., R.W., and D.A. contributed to data analysis, scientific presentation, discussion, and manuscript writing.

## Data and materials availability

All data needed to evaluate the conclusions in the paper are present in the paper or the Supplementary Materials.

## Supplementary Information

### Supplementary Figures

**Fig. S1.**
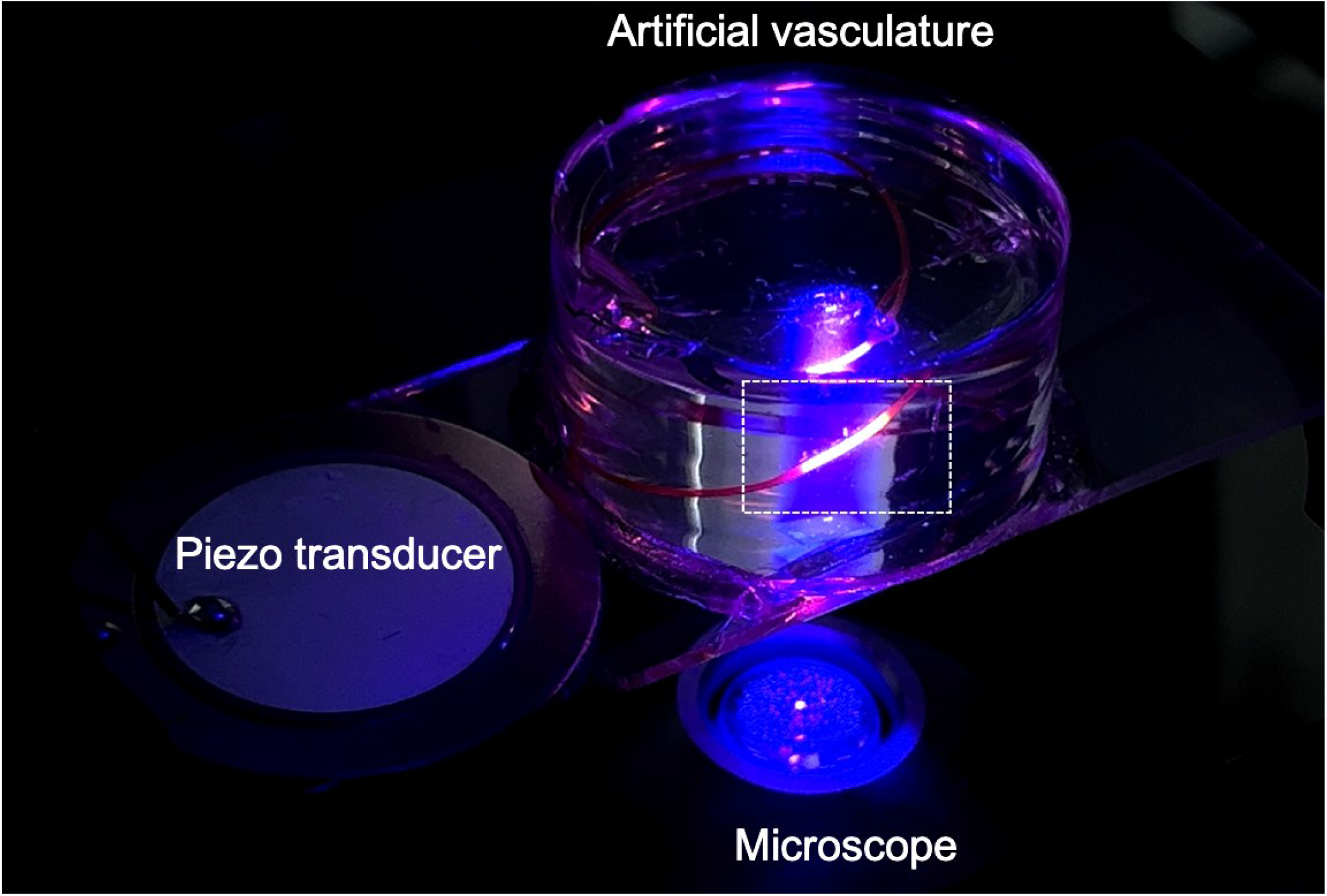
Experimental setup. The experimental setup consists of the acoustic excitation system and an acoustic manipulation chip. The acoustic manipulation chip is mounted on an inverted microscope and the movement of the microrobot is imaged by high-speed and high-sensitivity cameras.

**Fig. S2.**
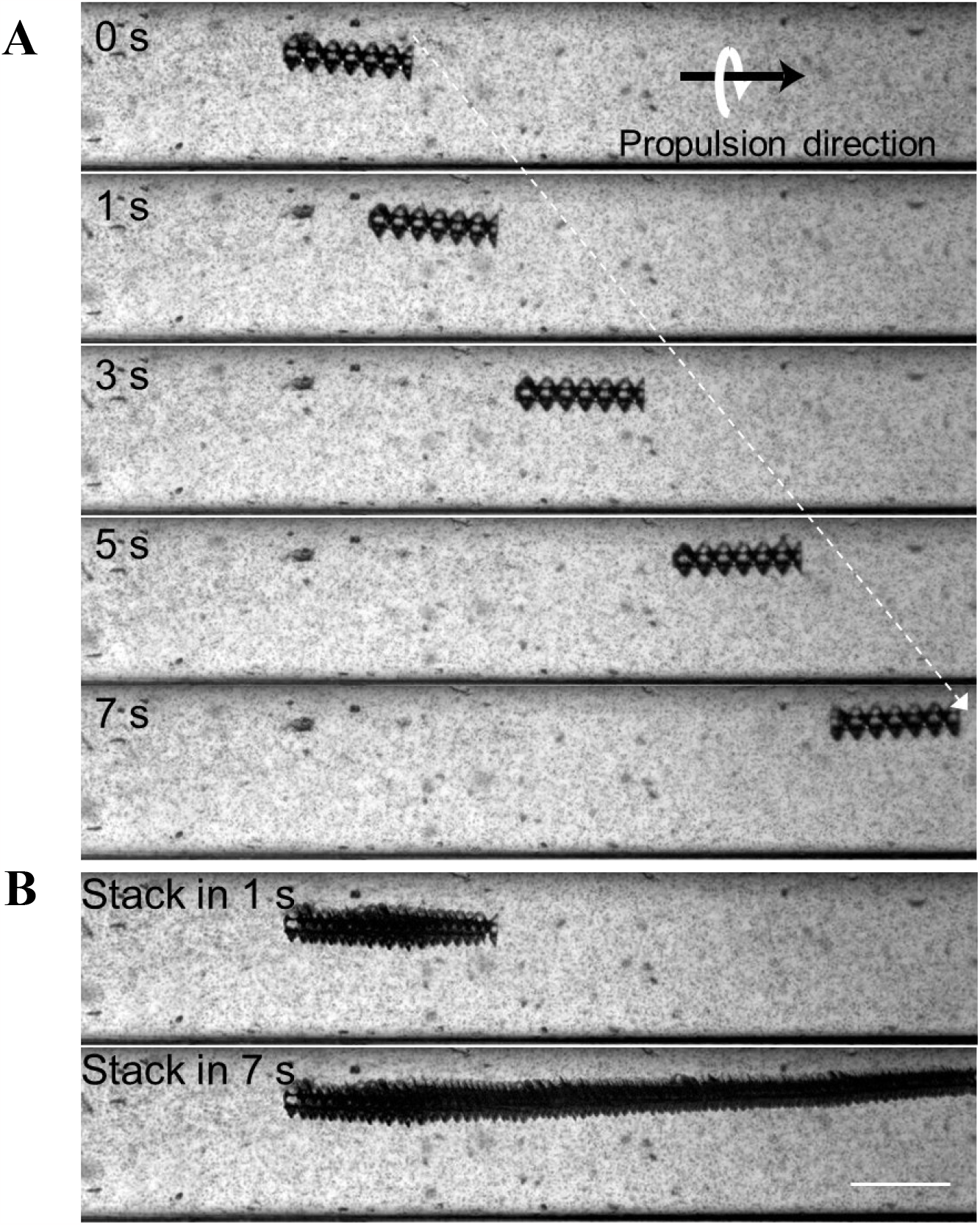
Streaming observation. (**A**) Image sequence with a moving microrobot and many small tracer particles, showing that there is no significant bulk flow that might drive the microrobot (see **Movie S2**). (**B**) Corresponding stacks of images with different time periods. Scale bar, 250 μm.

**Fig. S3.**
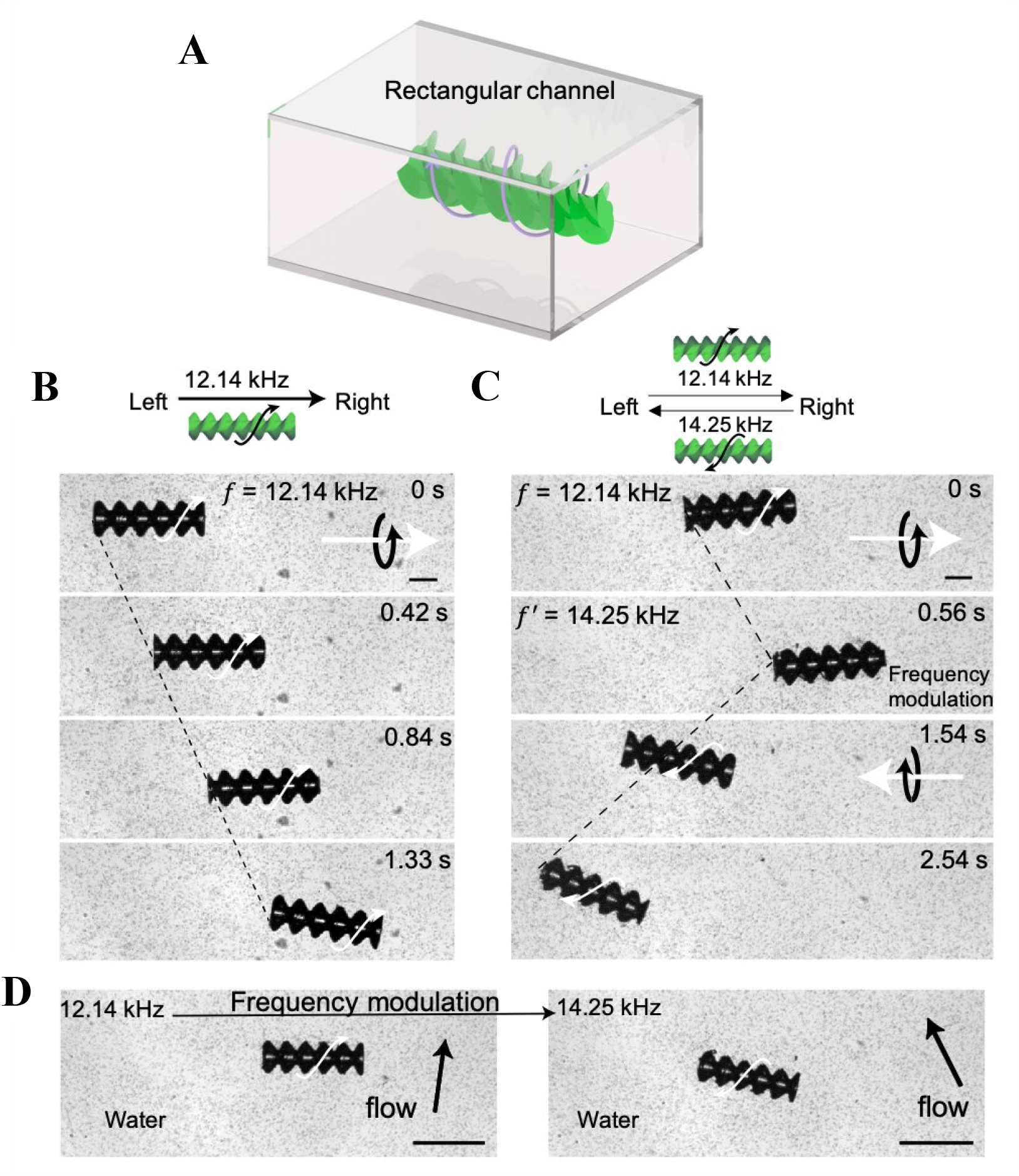
Manipulation of a microrobot in the rectangular chamber. (**A**) Setup for 2D manipulation of the microrobot in the chamber. (**B**) The microrobot is driven from left to right by acoustics (unidirectional motion). (**C**) The microrobot is actuated first from left to right to then from right to left by modulating the excitation frequency of the transducer (bidirectional motion). (**D**) The bulk flow is driven by acoustics in the rectangular channel when no microrobot is present. The microrobot does not follow the bulk flow, indicating the ability to move against flow in the chamber to some extent.

**Fig. S4.**
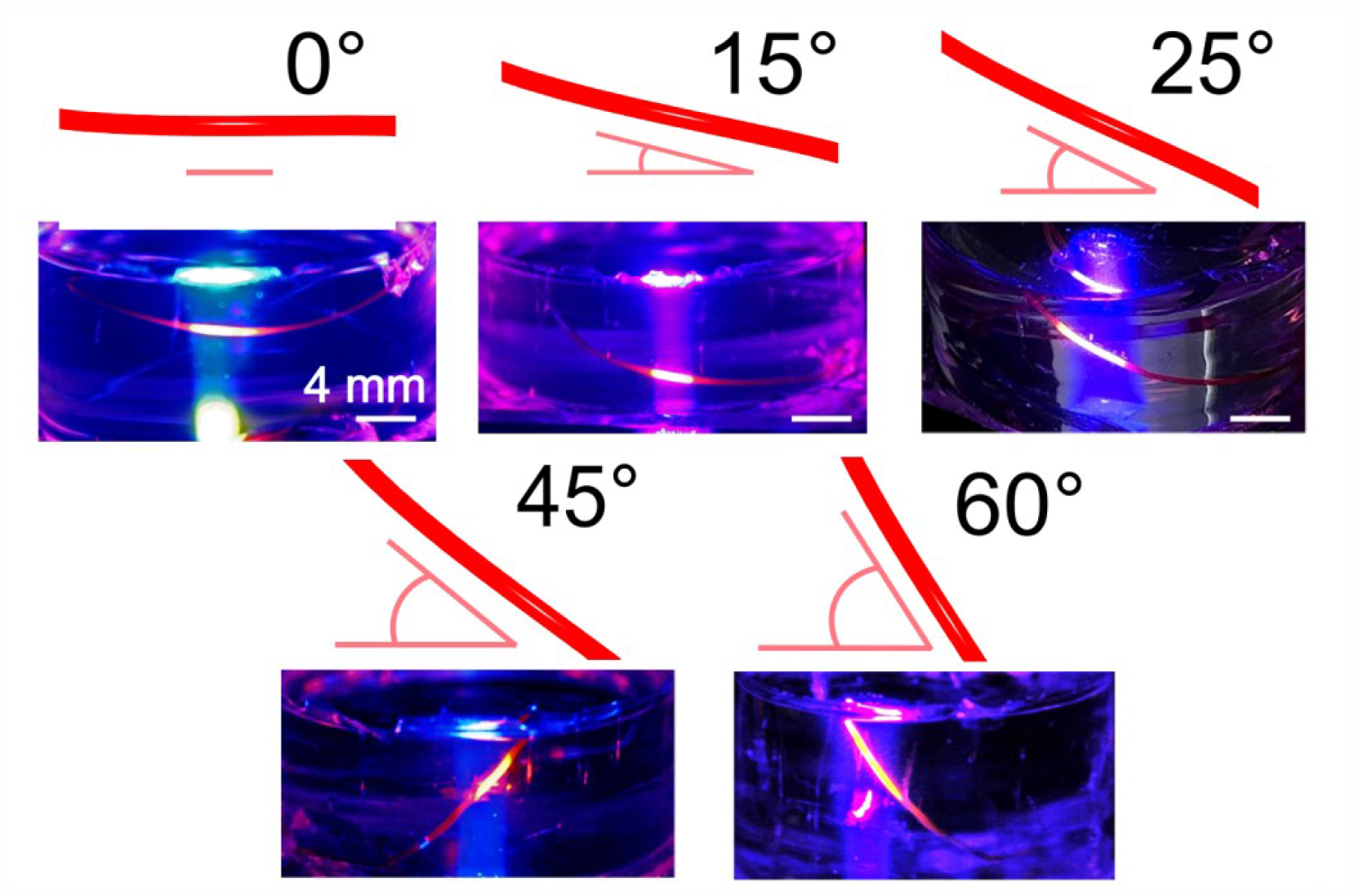
Artificial vasculatures created from PDMS with 0°, 15°, 25°, 45°, and 60° inclinations.

### Supplementary Note 1: Numerical simulations

Finite element computer simulations of the propulsion of the microrobot are performed using the commercial COMSOL Multiphysics software (v5.6, Burlington, MA). Three physical fields and the modules Thermoviscous acoustics, Laminar flow, and Solid mechanics are used.

#### Governing equations

We use the Navier-Stokes equations (see **Eqs. (1)** and **(2)** in the main text) together with the continuity equation (*1, 2*)

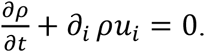

Here, *ρ* is the mass density of the fluid and *u*_*i*_ the *i*th element of the fluid’s velocity field. As is usual in acoustofluidics, a perturbation approach is applied to these equations, which permits the implementation of more efficient solvers, especially if we are only interested in a time-averaged result of the forces and torques acting on the microrobot. The first step is a Nyborg expansion *Σ* − *Σ*_0_ = *Σ*_1_ + *Σ*_2_+…, where the mass density, pressure (*p*), and velocity fields are expanded up to second order accuracy (*3, 4*):

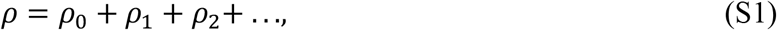

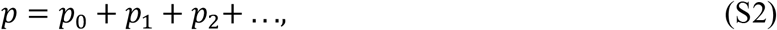

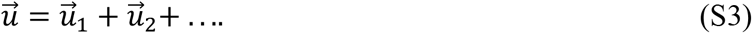

The subscripts 0, 1, and 2 denote different orders in the Nyborg expansion. Applying this expansion to the Navier-Stokes equations yields (*3, 4*)

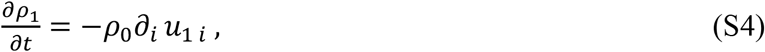

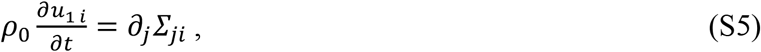

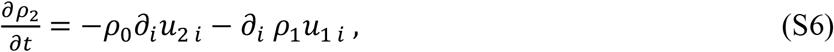

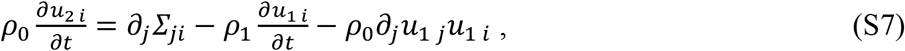

where the first- and second-order momentum-stress tensors are

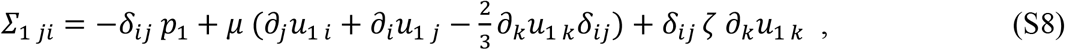

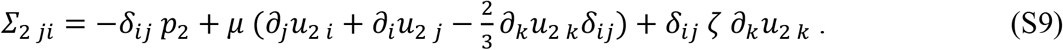

Since the first-order contribution has a harmonic time dependence, **Eqs. (S6)** and **(S7)** can be rewritten in time-averaged form (where ⟨⟩ denotes a time average over one wave period):

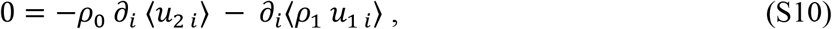

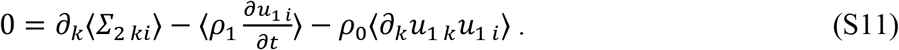

Hence, the first-order fields act as the source terms for second-order fields. The second-order terms can therefore be addressed by implementing the first-order terms in COMSOL. Applying the time-averaged momentum-stress tensor of the first and second order as shown in **Eqs. (4)** and **(5)** in the main text, the second-order total force 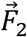 and torque 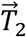 acting on the microrobot may be written as (*3, 5-8*)

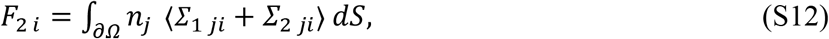

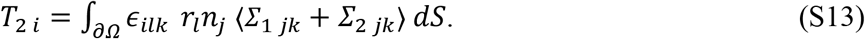

To get the radiation force 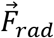 and torque 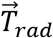 acting on the particle, a surface integral over the time-averaged second-order momentum-flux tensor ⟨*ρ*_0_*u*_1 *j*_*u*_1 *i*_⟩ may be used as an approximation for the acoustic streaming force and torque,

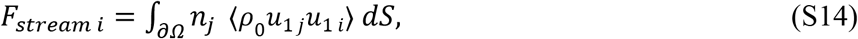

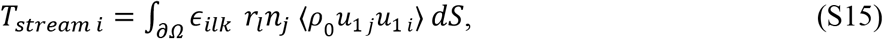

which are subtracted from the total force **(S12)** and torque **(S13)** to arrive at **Eqs. (6)** and **(7)** in the main text (*9*).

**Supplementary Table S1.**
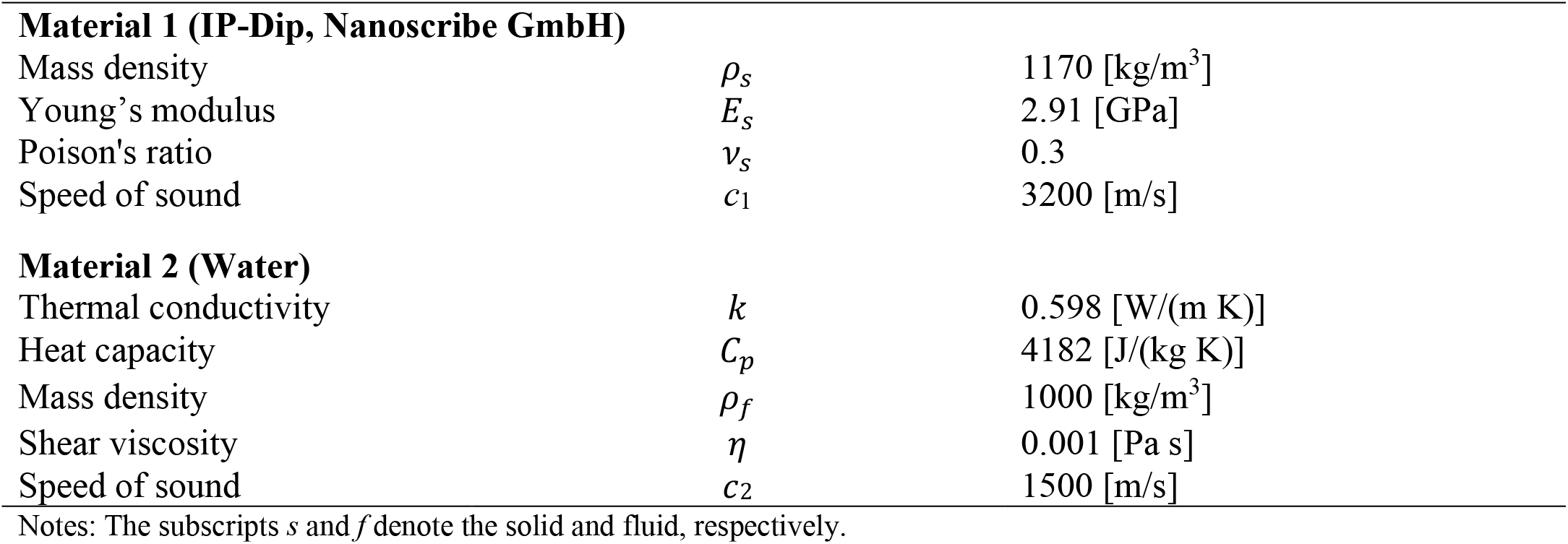
Parameters of materials used in the numerical simulations.

|  |  |  |
| --- | --- | --- |
| **Material 1 (IP-Dip, Nanoscribe GmbH)** |  |  |
| Mass density | $\rho_s$ | 1170 [kg/m<sup>3</sup>] |
| Young's modulus | $E_s$ | 2.91 [GPa] |
| Poisson's ratio | $\nu_s$ | 0.3 |
| Speed of sound | $c_1$ | 3200 [m/s] |
| **Material 2 (Water)** |  |  |
| Thermal conductivity | $k$ | 0.598 [W/(m K)] |
| Heat capacity | $C_p$ | 4182 [J/(kg K)] |
| Mass density | $\rho_f$ | 1000 [kg/m<sup>3</sup>] |
| Shear viscosity | $\eta$ | 0.001 [Pa s] |
| Speed of sound | $c_2$ | 1500 [m/s] |

Notes: The subscripts $s$ and $f$ denote the solid and fluid, respectively.

### Legends for Supplementary Movies

**Movie S1: Simulations of the microrobot**. The simulation model (see **Supplementary Note 1**) is analogous to the experimental configuration of the microrobot in the microchannel (the microrobot has diameter 100 μm and length 350 μm, the microchannel has diameter 500 μm). The acoustic excitation frequency is 13.5 kHz.

**Movie S2: Propulsion of the microrobot in 2D channel**. The motion of the microrobot in a microchannel is shown. There are no noticeable acoustic streaming and no acoustic standing wave in the microchannel. In contrast, the local bulk flow arising from the microrobot’s rotation is dominant. The acoustic excitation frequency and voltage are 13.5 kHz and 20 V_pp_, respectively. The video is recorded at ∼7 fps and played at 2x speed.

**Movie S3: Propulsion of the microrobot in a rectangular chamber**. The microrobot first rotates and swims in one direction at an acoustic excitation frequency of 12.15 kHz. Then, the frequency is tuned to 14.25 kHz and the microrobot immediately starts to rotate and swim in the opposite direction. The excitation voltage is 20 V_pp_. The video is recorded by a high-speed camera at 1070 fps and played at 214 fps (0.2x).

**Movie S4: 2D bidirectional propulsion of the microrobot in the artificial vasculature (circular-cross-section microfluidic channels)**. The microrobot is first propelled from left to right in the microchannel at an acoustic excitation frequency of 13.5 kHz. Then, the frequency is tuned to 18.5 kHz and the microrobot propels from right to left. The voltage of left-to-right propulsion is 20 V_pp_ and the voltage of right-to-left propulsion is 45 V_pp_. The video is recorded at ∼7 fps and played at speed 5x.

**Movie S5: Effects of the effective radius ratio** rd **on the translation and rotation velocities of the microrobot in the microchannel**. rd is defined as the ratio of the radius of the microrobot’s cylindrical core to the outer radius of the microrobot (100 μm). rd controls the surface area of the helical vanes (for rd=1 the m*i*crorobot *i*s a cyl*i*nder, for decreasing rd the surface area of the vanes increases). When reducing rd, the microrobot moves faster. Experiments were performed at 13.5 kHz and 20 V_pp_ in the microchannel of diameter 500 μm. The video is recorded at ∼7 fps and played at speed 2x.

**Movie S6: Effects of the reduced helical pitch** *h*_*d*_ = *P*/(2*R)* **on the microrobot’s propulsion**. Microrobots with 3.5, 7, and 14 turns, corresponding to a pitch of 216 μm, 108 μm, and 54 μm (i.e., *h*_*d*_=2.16, 1.08, 0.54), respectively, are shown. The microrobots’ outer diameter is kept constant at 100 μm, their length is kept at 350 μm and the microchannel had diameter 500 μm. Microrobots with larger *h*_*d*_ propel faster. The video is recorded at ∼7 fps and played at speed 2x.

**Movie S7: 3D manipulation of the microrobot in the microchannels with different inclinations (0°, 15°, 25°, 45°, 60°, and 75°)**. The inclination is defined as the angle between the inclined channel and the horizontal plane. Here, the microrobot is the same for each channel (diameter 500 µm). The video is record at ∼7 fps and play at speed 2x.

**Movie S8: 3D upward and downward manipulation of the microrobot in microchannels inclined at 15°**.

The video is recorded at ∼7 fps and played at speed 2x.

## REFERENCES AND NOTES

1. N. W. Charon, S. F. Goldstein, Genetics of motility and chemotaxis of a fascinating group of bacteria: The spirochetes. Annu. Rev. Genet. 36, 47–73 (2002).

2. C. Li, A. Motaleb, M. Sal, S. F. Goldstein, N. W. Charon, Spirochete periplasmic flagella and motility. J. Mol. Microbiol. Biotechnol. 2, 345–354 (2000).

3. A. Ghosh, D. Dasgupta, M. Pal, K. I. Morozov, A. M. Leshansky, A. Ghosh, Helical nanomachines as mobile viscometers. Adv. Funct. Mater. 28, 1705687 (2018).

4. L. Zhang, J. J. Abbott, L. Dong, B. E. Kratochvil, D. Bell, B. J. Nelson, Artificial bacterial flagella: Fabrication and magnetic control. Appl. Phys. Lett. 94, 064107 (2009).

5. X. Yan, Q. Zhou, M. Vincent, Y. Deng, J. Yu, J. Xu, T. Xu, T. Tang, L. Bian, Y.-X. J. Wang, K. Kostarelos, L. Zhang, Multifunctional biohybrid magnetite microrobots for imaging-guided therapy. Sci. Robot. 2, eaaq1155 (2017).

6. F. Zhang, Z. Li, Y. Duan, A. Abbas, R. Mundaca-Uribe, L. Y. Luan, W. Gao, R. H. Fang, L. Zhang, J. Wang, Gastrointestinal tract drug delivery using algae motors embedded in a degradable capsule. Sci. Robot. 7, eabo4160 (2022).

7. L. Soler, S. Sanchez, Catalytic nanomotors for environmental monitoring and water remediation. Nanoscale 6, 7175–7182 (2014).

8. S. K. Srivastava, M. Guix, O. G. Schmidt, Wastewater mediated activation of micromotors for efficient water cleaning. Nano Lett. 16, 817–821 (2016).

9. M. Liu, T. Zentgraf, Y. Liu, G. Bartal, X. Zhang, Light-driven nanoscale plasmonic motors. Nat. Nanotechnol. 5, 570–573 (2010).

10. S. Ghosh, F. Mohajerani, S. Son, D. Velegol, P. J. Butler, A. Sen, Motility of enzyme-powered vesicles. Nano Lett. 19, 6019–6026 (2019).

11. A. C. Hortelao, C. Simo, M. Guix, S. Guallar-Garrido, E. Julian, D. Vilela, L. Rejc, P. Ramos-Cabrer, U. Cossio, V. Gomez-Vallejo, T. Patino, J. Llop, S. Sanchez, Swarming behavior and in vivo monitoring of enzymatic nanomotors within the bladder. Sci. Robot. 6, eabd2823 (2021).

12. T. Patino, A. Porchetta, A. Jannasch, A. Lladó, T. Stumpp, E. Schäffer, F. Ricci, S. Sánchez, Self-sensing enzyme-powered micromotors equipped with pH-responsive DNA nanoswitches. Nano Lett. 19, 3440–3447 (2019).

13. K. Villa, H. Sopha, J. Zelenka, M. Motola, L. Dekanovsky, D. C. Beketova, J. M. Macak, T. Ruml, M. Pumera, Enzyme-photocatalyst tandem microrobot powered by urea for Escherichia coli biofilm eradication. Small 18, 2106612 (2022).

14. G. Loget, A. Kuhn, Electric field-induced chemical locomotion of conducting objects. Nat. Commun. 2, 535 (2011).

15. W. Gao, S. Sattayasamitsathit, K. M. Manesh, D. Weihs, J. Wang, Magnetically powered flexible metal nanowire motors. J. Am. Chem. Soc. 132, 14403–14405 (2010).

16. H. Gu, E. Hanedan, Q. Boehler, T.-Y. Huang, A. J. T. M. Mathijssen, B. J. Nelson, Artificial microtubules for rapid and collective transport of magnetic microcargoes. Nat. Mach. Intell. 4, 678–684 (2022).

17. Y. Kim, X. Zhao, Magnetic soft materials and robots. Chem. Rev. 122, 5317–5364 (2022).

18. I. C. Yasa, H. Ceylan, U. Bozuyuk, A.-M. Wild, M. Sitti, Elucidating the interaction dynamics between microswimmer body and immune system for medical microrobots. Sci. Robot. 5, eaaz3867 (2020).

19. H. Zhou, C. C. Mayorga-Martinez, S. Pane, L. Zhang, M. Pumera, Magnetically driven micro and nanorobots. Chem. Rev. 121, 4999–5041 (2021).

20. A. Aghakhani, A. Pena-Francesch, U. Bozuyuk, H. Cetin, P. Wrede, M. Sitti, High shear rate propulsion of acoustic microrobots in complex biological fluids. Sci. Adv. 8, eabm5126 (2022).

21. D. Ahmed, T. Baasch, N. Blondel, N. Laubli, J. Dual, B. J. Nelson, Neutrophil-inspired propulsion in a combined acoustic and magnetic field. Nat. Commun. 8, 770 (2017).

22. D. Ahmed, C. Dillinger, A. Hong, B. J. Nelson, Artificial acousto-magnetic soft microswimmers. Adv. Mat. Tech. 2, 1700050 (2017).

23. D. Ahmed, A. Sukhov, D. Hauri, D. Rodrigue, M. Gian, J. Harting, B. Nelson, Bio-inspired acousto-magnetic microswarm robots with upstream motility. Nat. Mach. Intell. 3, 116–124 (2021).

24. C. Dillinger, N. Nama, D. Ahmed, Ultrasound-activated ciliary bands for microrobotic systems inspired by starfish. Nat. Commun. 12, 6455 (2021).

25. D. Ahmed, T. Baasch, B. Jang, S. Pane, J. Dual, B. J. Nelson, Artificial swimmers propelled by acoustically activated flagella. Nano Lett. 16, 4968–4974 (2016).

26. C. Hong, Z. Ren, C. Wang, M. Li, Y. Wu, D. Tang, W. Hu, M. Sitti, Magnetically actuated gearbox for the wireless control of millimeter-scale robots. Sci. Robot. 7, eabo4401 (2022).

27. P. E. Dupont, B. J. Nelson, M. Goldfarb, B. Hannaford, A. Menciassi, M. K. O’Malley, N. Simaan, P. Valdastri, G.-Z. Yang, A decade retrospective of medical robotics research from 2010 to 2020. Sci. Robot. 6, eabi8017 (2021).

28. P. Fischer, B. J. Nelson, Tiny robots make big advances. Sci. Robot. 6, eabh3168 (2021).

29. E. W. H. Jager, O. Inganas, I. Lundstrom, Microrobots for micrometer-size objects in aqueous media: Potential tools for single-cell manipulation. Science 288, 2335–2338 (2000).

30. W. Wang, L. A. Castro, M. Hoyos, T. E. Mallouk, Autonomous motion of metallic microrods propelled by ultrasound. ACS Nano 6, 6122–6132 (2012).

31. L. Ren, N. Nama, J. M. McNeill, F. Soto, Z. Yan, W. Liu, W. Wang, J. Wang, T. E. Mallouk, 3D steerable, acoustically powered microswimmers for single-particle manipulation. Sci. Adv. 5, eaax3084 (2019).

32. A. Aghakhani, O. Yasa, P. Wrede, M. Sitti, Acoustically powered surface-slipping mobile microrobots. Proc. Natl. Acad. Sci. U.S.A. 117, 3469–3477 (2020).

33. R. C. Johnson, F. W. Hyde, C. M. Rumpel, Taxonomy of the lyme disease spirochetes. Yale. J. Biol. Med. 57, 529–537 (1984).

34. W. L. M. Nyborg, Acoustic streaming. In Physical acoustics. (Academic Press, Vermont, 1965), vol. 2, pp. 265–331.

35. J. Voß, R. Wittkowski, On the shape-dependent propulsion of nano- and microparticles by traveling ultrasound waves. Nanoscale Adv. 2, 3890–3899 (2020).

36. J. Voss, R. Wittkowski, Orientation-dependent propulsion of triangular nano- and microparticles by a traveling ultrasound wave. ACS Nano 16, 3604–3612 (2022).

37. J. Dual, P. Hahn, I. Leibacher, D. Moller, T. Schwarz, J. Wang, Acoustofluidics 19: Ultrasonic microrobotics in cavities: Devices and numerical simulation. Lab Chip 12, 4010–4021 (2012).

38. J. T. Karlsen, H. Bruus, Forces acting on a small particle in an acoustical field in a thermoviscous fluid. Phys. Rev. E 92, 043010 (2015).

39. E. B. Lima, J. P. Leão-Neto, A. S. Marques, G. C. Silva, J. H. Lopes, G. T. Silva, Nonlinear interaction of acoustic waves with a spheroidal particle: Radiation force and torque effects. Phys. Rev. Appl. 13, 064048 (2020).

40. P. B. Muller, H. Bruus, Theoretical aspects of microchannel acoustofluidics: Thermoviscous corrections to the radiation force and streaming. Procedia IUTAM 10, 410–415 (2014).

41. H. Bruus, Acoustofluidics 2: Perturbation theory and ultrasound resonance modes. Lab Chip 12, 20–28 (2012).

42. M. Settnes, H. Bruus, Forces acting on a small particle in an acoustical field in a viscous fluid. Phys. Rev. E 85, 016327 (2012).

43. J. Voß, J. Jeggle, R. Wittkowski, HydResMat–FEM-based code for calculating the hydrodynamic resistance matrix of an arbitrarily-shaped colloidal particle. GitHub: HV59/HydResMat. Zenodo. doi: 10.5281/zenodo.3541588 (2019).

44. J. Happel, H. Brenner, Low Reynolds number hydrodynamics: With special applications to particulate media. (Springer Science & Business Media, The Hague, 1983), vol. 1.

45. D. Ahmed, M. Lu, A. Nourhani, P. E. Lammert, Z. Stratton, H. S. Muddana, V. H. Crespi, T. J. Huang, Selectively manipulable acoustic-powered microswimmers. Sci. Rep. 5, 9744 (2015).

46. S. Lee, J. Y. Kim, J. Kim, A. K. Hoshiar, J. Park, S. Lee, J. Kim, S. Pane, B. J. Nelson, H. Choi, A needle-type microrobot for targeted drug delivery by affixing to a microtissue. Adv. Healthc. Mater. 9, 1901697 (2020).

47. J. Li, S. Sattayasamitsathit, R. Dong, W. Gao, R. Tam, X. Feng, S. Ai, J. Wang, Template electrosynthesis of tailored-made helical nanoswimmers. Nanoscale 6, 9415–9420 (2014).

48. C. Peters, O. Ergeneman, P. D. W. García, M. Müller, S. Pané, B. J. Nelson, C. Hierold, Superparamagnetic twist-type actuators with shape-independent magnetic properties and surface functionalization for advanced biomedical applications. Adv. Funct. Mater. 24, 5269–5276 (2014).

49. L. Schwarz, D. D. Karnaushenko, F. Hebenstreit, R. Naumann, O. G. Schmidt, M. Medina-Sanchez, A rotating spiral micromotor for noninvasive zygote transfer. Adv. Sci. 7, 2000843 (2020).

50. J. Lei, P. Glynne-Jones, M. Hill, Comparing methods for the modelling of boundary-driven streaming in acoustofluidic devices. Microfluid. Nanofluid. 21, 23 (2017).

51. H. Bruus, Acoustofluidics 1: Governing equations in microfluidics. Lab Chip 11, 3742–3751 (2011).

52. H. Bruus, Acoustofluidics 10: Scaling laws in acoustophoresis. Lab Chip 12, 1578–1586 (2012).

